# An electrophysiological study about the pharmacological manipulation of the immediate consequences of a spinal trauma reveals a crucial role for TRPV4 antagonism

**DOI:** 10.1101/2024.10.03.616499

**Authors:** Atiyeh Mohammadshirazi, Giuliano Taccola

## Abstract

A physical trauma to the spinal cord produces an immediate massive depolarizing injury potential accompanied both by a transient episode of spinal hypoxia, and an extensive cell loss at the level of injury, which interrupts conduction of longitudinal input along white matter tracts. Afterwards, the transient hypotonia and areflexia characterize the following spinal shock phase. The relationship between the extent of injury potentials and spinal cord injury (SCI) progression, as well as the potential pharmacological modulation of the immediate consequences of a trauma, have not yet been explored. To limit the peak of injury potentials and speed up recovery of reflex motor responses, we serially applied selective neurochemicals in the exact moment of an experimental physical trauma delivered through a calibrated device impacting the mid-thoracic cord of an entire CNS preparation of neonatal rats. Continuous lumbar root recordings monitored baseline DC-levels and reflex responses elicited by trains of electric pulses applied to sacrocaudal afferents. In uninjured preparations, each agent showed distinct effects on baseline polarization, modulation of synaptic responses, and appearance of bursting activity. Interestingly, neurochemicals acting on glutamatergic-, adenosinergic-, glycinergic- or GABAergic receptors, did not affect the monitored outcome when each parameter was normalized against pre-injury values. Conversely, the selective TRPV4 antagonist, RN1734, unlike the TRPA1 antagonist, AP18, reduced peak of injury potentials and speeded up full recovery of reflex responses within 1 min from trauma. Similarly, blockage of gap junctions quickly, yet partially, restored motor reflexes, while antagonism of GABA_A_ receptors restored full reflexes, though slightly later. The current study indicates that both mechanosensitive TRPV4 receptors and GABAergic transmission reduce the immediate pathological consequences of a trauma when applied at the moment of impact, envisaging a clinical translation for preventing accidental spinal lesions during the most delicate spinal surgeries.

## Introduction

We have recently described the immediate (150-200 ms) functional changes occurring when combining a physical trauma applied to the spinal cord of the entire CNS isolated from neonatal rats, with the use of an ad hoc designed micro impactor (Mohammadshirazi et al., 2023, 2024). Each impact consistently triggered an early depolarization that paralleled the injury potential reported in preclinical studies, with an amplitude that was proportional to the strike intensity and modulated by varying extracellular ion concentrations. At injury level, extensive cellular loss and disconnection of input along the spinal cord parallel a severe clinical spinal cord injury (SCI). Moreover, hypotonia and areflexia that characterize clinical spinal shocks were mimicked in vitro by the suppression of spontaneous motor activity from all VRs and by the transient failure of afferent pulses, which reflect the impairment of spinal synaptic transmission. In the present study, we explored the possibility to limit the extent of depolarization after injury, promptly restore reflexes and protect neuronal networks, through the pharmacological manipulation of membrane receptors mediating fast ionic currents in the spinal cord. Besides the multiple neurotransmitter systems investigated so far to reactivate sensory motor networks after injury (Musienko et al., 2011), neurochemicals acting on ionotropic receptors implicated in the pathophysiology of acute SCIs, might represent an optimal target to limit the sudden depolarization following an impact to the spinal cord. In particular, since low extracellular concentrations of chloride ions reduce the peak of injury potentials (Mohammadshirazi et al., 2024), chloride-mediated fast inhibitory neurotransmission can be exploited by acting on both, glycine and GABA_A_ receptors. Furthermore, a massive surge of glutamate has been reported as an early consequence of a SCI (Liu et al., 1991; McAdoo et al., 1999, 2000; Xu, 2004), suggesting to reduce the early functional changes induced by the trauma by blocking ionotropic glutamate receptors already during the impact, using APV and CNQX, which mostly suppress spinal synaptic transmission (Mayer and Westbrook, 1987; Bracci et al., 1996a). However, apart from ionotropic receptors, the wave of ions sustaining tissue depolarization could also spread along the cord through gap junctions. Gap junctions have been linked to axonal dysfunctions after SCI, as their blockage through carbenoxolone limited the loss of impulse conduction (Goncharenko et al., 2014).

Micro dialysis sampling at injury level also highlighted the acute release of adenosine (McAdoo et al., 2000), suggesting to limit the first consequences of a trauma by applying either adenosine, or the broad antagonist of adenosine receptors, caffeine, or more selectively, the A1 receptor blocker, DPCPX, which has been reported to modulate sensorimotor networks in the spinal cord (McAdoo et al., 2000; Taccola et al., 2020).

Due to stretch forces produced on the membrane of spinal neurons at the injury epicenter, mechanoreceptors might also be involved in the trauma. Mechanosensitive neurons express a broad class of mechanically gated ion channels, which are activated by sheer forces applied to close-by membranes and are coupled to cations currents (Garcia-Elias et al., 2014). However, although described in the spinal cord of lampreys (Grillner et al., 1982) and lizards (Alibardi, 2019), only few evidence exists about the presence of functional mechanoreceptors in the ventral spinal cord of mammals.

Among the four superfamilies of mechanoreceptors (Valle et al., 2012), the TRP family includes the transient receptor potential, vanilloid 4 (TRPV4). TRPV4 is a calcium-permeable non-selective cation channel expressed by astrocytes and neurons, and is widespread in the central nervous system (CNS), including the spinal cord, although at lower levels (Kumar et al., 2020). Once activated, TRPV4 depolarizes cell membrane, but also modulates ligand-gated chloride channels, such as GABA_A_ receptors and strychnine-sensitive GlyR (Qi et al., 2018). TRPV4 channels are activated by a wide range of stimuli, as they are osmoreceptors sensing mechanical stimulation due to cell swelling, but also thermosensors for warm temperatures above 27° C (Kumar and Han, 2022). Impacts to the spinal cord increase the expression of TRPV4 during the early inflammatory phase of an SCI, proportionately to the severity of trauma (Kumar et al., 2020; Kumar and Han, 2022). Genetic suppression of TRPV4 or intraperitoneal administration of the selective pharmacological TRPV4 antagonist 1 h after SCI, enhanced neuroprotection against SCI-induced endothelial damage with preserved blood-spinal cord barrier integrity, attenuated neuroinflammation, and reduced glial scarring at the epicenter of injury, with some motor recovery in hindlimbs (Kumar et al., 2020).

Our hypothesis is that mechanoreceptors are immediately activated by an impact and, even before the opening of the ionotropic receptors considered herein, they could be responsible for the extracellular ion imbalance that sustains both injury potentials and transient reflex suppression during spinal shock. To this aim, in the present study, spinal impacts of equal severity were singularly performed on different preparations. Each preparation was then perfused with a single neuroactive drug before the impact and for the following recovery phase. Amplitude and latency of injury potentials and time-courses of recovery of spinal reflexes were compared to untreated injured cords. Surprisingly, glutamatergic-, adenosinergic-, glycinergic- or GABAergic receptors, were not implicated in sustaining the peak of injury potentials that follow a physical trauma to the spinal cord. However, the selective TRPV4 antagonist, RN1734, but not so the TRPA1 antagonist, AP18, limited the insurgence of the immediate consequences of the impact when applied at the moment of compression. Moreover, blockage of gap junctions speeded up the partial recovery of reflex responses, while the antagonism of GABA_A_ receptors fully restored reflexes, though slightly later. This evidence shows the crucial roles played by mechanosensitive TRPV4 receptors, as well as by GABAergic transmission, in the pathophysiology of an acute spinal trauma, suggesting potential implications for a clinical translation aimed at limiting accidental damages to the cord during the most delicate spinal surgeries.

## Methods

To explore the pharmachological modulation of the immediate consequences of a traumatic injury to spinal cord, we adopted the in vitro preparation of the entire CNS isolated from neonatal rats (Mohammadshirazi et al., 2023; Apicella and Taccola, 2023) with intact dorsal laminae (Mohammadshirazi et al., 2024). Continuous nerve recordings were performed from VRrL5, while sacrocaudal afferents were electrically stimulated at low frequency throughout the experiment (Fig. 1A). A calibrated severe impact was provided to the ventral side of the tenth segment of the thoracic spinal cord (T10).

**Figure 1.**
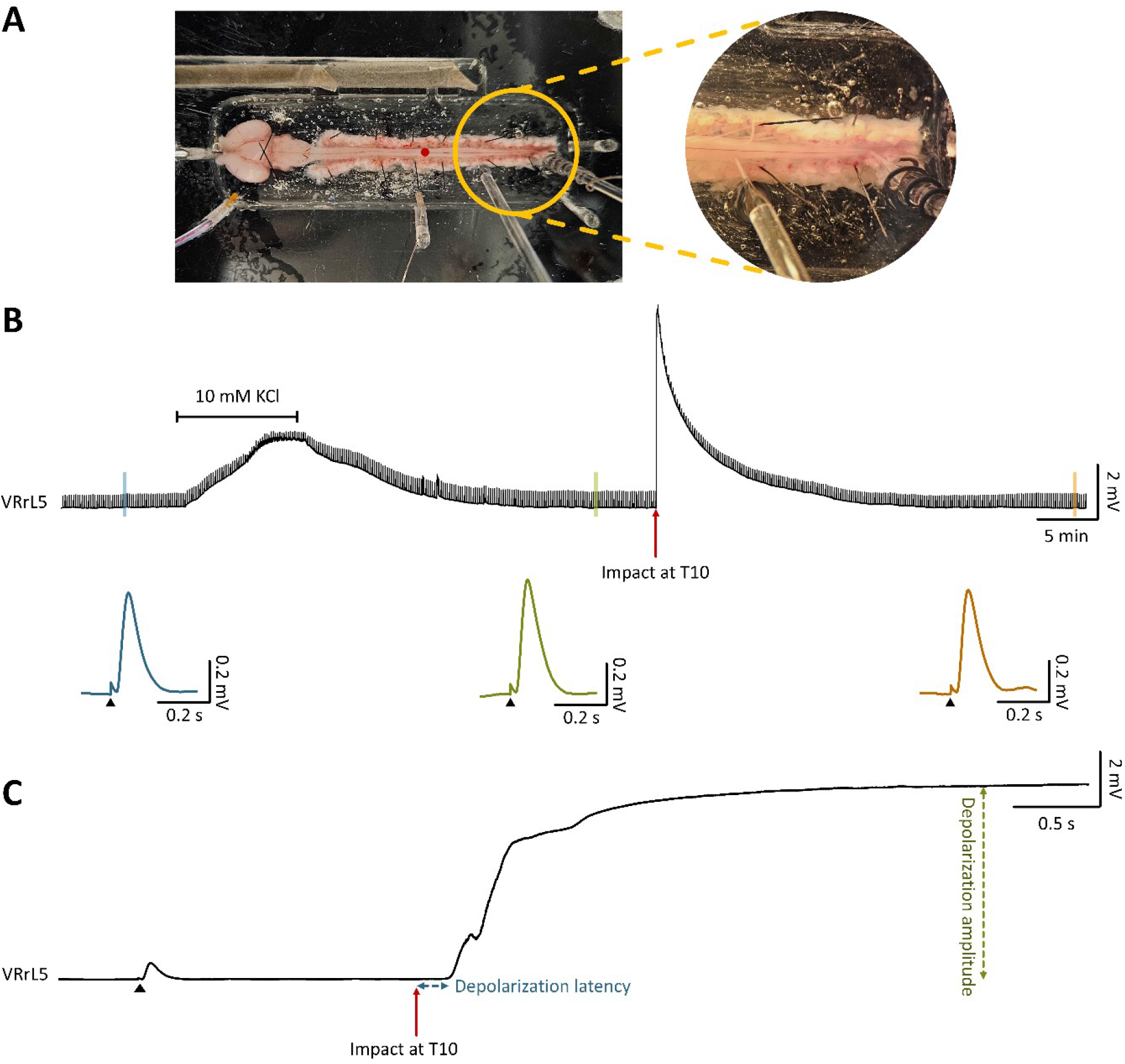
Transient depolarization induced by a physical injury to the spinal cord in an in vitro CNS preparation. **A**. A picture of the electrophysiological setup for in vitro preparations of the entire CNS with intact dorsal aspects of vertebrae. Throughout the experiment, continuous recordings were derived from VRrL5, as electrical stimulation was applied to sacrocaudal afferents every 10 seconds. Physical impact was performed at T10 (red dot). **B**. A long continuous recording from VRrL5, while the cord was impacted at T10 (red arrow). Ten minutes perfusion with 10 mM KCl represent an internal control to quantify the high recruitment of motor pools. Below, three examples of electrically induced reflex responses are magnified before (blue) and after (green) KCl application, and once recovered from impact-induced depolarization (orange). **C**. A magnified 6-second trace extracted from B, highlights the instants around the impact and defines latency (blue dotted arrow) and amplitude (green dotted arrow) of impact-induced depolarization.

### In vitro preparation of the isolated entire CNS

All procedures were approved by the International School for Advanced Studies (SISSA) ethics committee and are in accordance with the guidelines of the National Institutes of Health (NIH) and the Italian Animal Welfare Act 24/3/2014 n. 26, implementing the European Union directive on animal experimentation (2010/63/EU). Every measure was taken to minimize the number of animals used and to ensure their welfare.

In vitro preparations of the entire isolated central nervous system (CNS; Mohammadshirazi et al., 2023; Apicella and Taccola, 2023) coming from 94 postnatal (P0 - P2.5) Wistar rats were utilized in this study. Following cryoanesthesia (Phifer and Terry, 1986; Danneman and Mandrell, 1997; Goldberg, 2015) and the disappearance of the paw pinch reflex, quick surgical procedures were performed including removal of: forehead at orbital line, rib cage, internal organs, and forelimbs. Dissection continued under the microscope in a petri dish filled with an oxygenated Krebs solution that contained (in mM): 113 NaCl, 4.5 KCl, 1 MgCl_2_7H_2_O, 2 CaCl_2_, 1 NaH_2_PO_4_, 25 NaHCO_3_, and 30 glucose, gassed with 95% O_2_ - 5% CO_2_, pH 7.4, 299.62 ± 3.2 mOsm/kg. Craniotomy and ventral laminectomy were then performed, keeping dorsal vertebra and dorsal root ganglia (DRG) intact (Mohammadshirazi et al., 2024), and the preparation was maintained in the oxygenated Krebs solution at room temperature for 15 minutes and thus transferred to the recording chamber with perfusing oxygenated Krebs solution (7 mL/min) and controlled temperature of 25-27° C (TC-324C Warner Instruments, USA). For electrophysiological recordings, the preparation was fixed ventral side up and selected VRs were detached from DRGs.

### Extracellular recordings

Monopolar suction electrodes were created by pulling tight-fitting glass pipettes (1.5 mm outer diameter, 0.225 mm wall thickness; Hilgenberg, Germany) and used to obtain DC-coupled recordings from VRrL5. Electrodes were connected to a differential amplifier (DP-304, Warner Instruments, Hamden, CT, USA) and signals were acquired with X 1000 gain, 0.1 Hz high-pass and 10 kHz low-pass filter. After noise elimination of analog signals (D400, Digitimer Ltd, UK), traces were digitized with a sampling rate of 5 kHz and low-pass filtrated at 10 Hz (Digidata 1440, Molecular Devices Corporation, Downingtown, PA, USA; digital Bessel) and visualized real-time (software Clampex 10.7, Molecular Devices Corporation, Downingtown, PA, USA).

### Electrical Stimulation

A programmable stimulator (STG4002, Multichannel System, Reutlingen, Germany) and bipolar glass suction electrodes connected to two paired silver wires (500-300 µm) were utilized to produce electrical stimulations. Trains of rectangular electrical pulses of 40-160 µA intensity, 0.1 ms pulse duration, and 0.1 Hz frequency were supplied to the *cauda equina* (Etlin et al., 2010). Stimulus intensity was imputed as times to threshold (Th), which is defined as the lowest intensity required to elicit a slight deflection of VRrL5 baseline.

### Spinal cord injury

A custom-made and shielded micro-impactor device was used to provide a traumatic damage to the spinal cord of the entire CNS in vitro (Mohammadshirazi et al., 2024).

A calibrated impact was applied to the ventral surface of thoracic 10 (T10) while electrophysiological recordings were simultaneously performed. A dedicated software controlled the vertical movement of the impactor tip (diameter = 2 mm). To provide a severe damage to the spinal cord, the impactor tip descended quickly (time = 650 ms) into the cord by 2656 µm from the spinal surface (speed = 4 mm/s, acceleration and deceleration = 6.1 ± 0.05 mm/s) and promptly returned to the original position with the same acceleration and deceleration speed.

### Pharmacology

Based on their solubility product constant (Ksp), powders of pharmacological agents were dissolved in proper solvent to make concentrated stocks that were then serially diluted in oxygenated Krebs solution to reach the final concentration. All pharmacological agents were perfused into the recording chamber while the impact was being performed on the spinal cord. Concentration and time of application are mentioned in Table 1. All chemicals were provided by Sigma-Aldrich (Merk Life Science S.r.l., Italy).

**Table 1.** Pharmacological agents applied in the study, final concentration, type of solvent, and time of perfusion.

| Agents | Concentration (μM) | Solvent | Time of application before impact (min) | Time of application after impact (min) |
| --- | --- | --- | --- | --- |
| DL-2-amino-5- phosphonopentanoic acid (APV) | 100 | Water | 20 | 5 |
| 6-Cyano-7-nitroquinoxaline-2,3-dione disodium salt hydrate (CNQX) | 20 | DMSO | 20 | 5 |
| carbenoxolone disodium (CBX) | 100 | Water | 50 | 5 |
| adenosine (ADO) | 1000 | Krebs | 5 | 15 |
| caffeine | 300 | Water | 15 | 5 |
| 8-cyclopentyl-1,3-dipropylxanthine (DPCPX) | 5 | Sodium hydroxide | 5 | 5 |
| AP-18 | 20, 50, 66 | DMSO | 15 | 5 |
| RN-1734 | 10, 20, 50, 100 | DMSO | 15 | 5 |
| glycine (Gly) | 500 | Krebs | 20 | 5 |
| 1(S),9(R)-(-)-bicuculline methiodide (bic) | 20 | Water | 20 | 10 |
| strychnine (str) | 1 | Water | 20 | 10 |

### Data and statistical analysis

Data analysis was pursued using Clampfit 10.7 software (Molecular Devices Corporation, PA, USA), and statistical analysis was performed with GraphPad InStat 3.10 (Inc., San Diego, California, USA). The number of animals is indicated as “n”, and data is denoted as mean ± standard deviation (SD). The latency is defined as the time spanning from either, the electrical stimulation or the mechanical artifact, to the onset of increased DC level. Time to peak refers to the temporal delay from the artifact of stimulation to the peak of the subsequent evoked response. Power spectrum analysis was done to generate a frequency domain representation, revealing the power levels of different frequency components in the signal. Bursting activity was quantified in terms of power spectrum magnitude, expressed as Root Mean Square (RMS; Deumens et al., 2013) and measured with Clampex 10.7 (Molecular Devices Corporation, Downingtown, PA, USA). This statistical tool quantifies any increase in frequency and/or amplitude of rhythmic activity, defined as a complex rhythm composed of multiple harmonics. Prior to any other statistical tests, a normality test was performed. Parametric paired student t-test or non-parametric Wilcoxon matched-pairs test, or Kruskal-Wallis test following Dunn’s multiple comparisons to control data were utilized. Differences with a P value ≤ 0.05 are considered significant.

## Results

### A traumatic injury to the spinal cord generates a large, transient injury potential

In an exemplar experiment, the impact caused a depolarization that started after only 186.44 ms, peaked at 7.14 mV after 5.42 s from the impact (Fig. 1B, C), and recovered within approximately 25 minutes. In 12 preparations, the same traumatic injury generated a sudden potential with a latency of 188.62 ± 8.70 ms that reached a maximal mean amplitude of 5.50 ± 2.84 mV, and recovered to baseline in 11.99 ± 8.93 min.

A reference of the extent of depolarization that corresponds to a massive recruitment of motor pools in control was calculated for each animal. A ten-minute perfusion with a high K^+^ physiological solution (10 mM) depolarized VRs by 2.57 ± 1.09 mV (n = 12). In the same experiments, the mean amplitude of the depolarization induced by injury in physiological solution was more than double (226.79 ± 121.63 %, n = 12) than the one elicited by high potassium.

In a sample experiment, a train of electrical pulses (0.1 ms pulse duration, 0.1 Hz frequency, 160 µA intensity) was continuously applied to sacrocaudal afferents (Fig. 1B) and induced, in control, a motor reflex response of 0.49 mV amplitude, which diminished at the peak of the potassium-induced depolarization (29.92 % of control) with a full recovery (101.12 % of control) in 2.70 min after injury. Contrarywise, motor reflex responses totally disappeared at the peak of the injury-induced potential, with a substantial recovery (80.79 % of control; Fig. 1B) after 36.33 min from injury. Notewhorty, recovery of evoked responses after both, chemically and mechanically induced depolarizations, confirms that neither a perfusion with 10 mM KCl nor a severe impact at T10 reduced functionality of lumbar motor pools (Fig 1B). In 6 experiments, trains of stimuli (60 - 160 µA intensity) evoked mean motor reflex responses of 0.82 ± 0.25 mV amplitude in control, which were largely reduced at the peak of the potassium-induced depolarization (0.21 ± 0.19 mV) with a full recovery (96.86 ± 2.86 %) after 7.76 ± 3.68 min of washing. Moreover, mean reflexes were completely suppressed at the peak of the injury-induced potential, with a substantial recovery to baseline values after 18.49 ± 10.21 min from injury (n = 6).

### Effects of traumatic injury over baseline activity and electrically evoked responses

To modulate the extent of depolarization and the suppression of reflex responses after injury, in separate experiments, selected neurochemicals were perfused (Table 1) and their effects on baseline activity was explored before injury. Antagonism of inotropic glutamate receptors, using APV (100 µM) + CNQX (20 µM), slightly polarized DC level, reaching a peak of 0.29 ± 0.10 mV (n = 5) after 7.45 ± 2.20 min perfusion and remaining stable thereafter. With APV + CNQX, reflex responses gradually decreased, reaching 20 % of their original value in amplitude within 5.39 ± 1.25 minutes, and completely disappeared after 8.31 ± 1.77 minutes (n = 5). Adenosine (1 mM) polarized baseline by 0.34 ± 0.12 mV after 4.21 ± 0.59 minutes (n = 6), but also reduced and delayed reflex responses (Table 2, amplitude: P = 0.003, time to peak: P = 0.041, paired t-test). Caffeine (300 µM), a non-specific adenosine receptor antagonist, did not affect DC levels, but increased the latency of reflex responses (Table 2, P = 0.039, paired t-test). Following glycine (500 µM) application, a small depolarization of 1.70 ± 0.66 mV (n = 6) occurred within 2.31 ± 0.42 minutes. Within 6.06 ± 2.46 minutes, a spontaneous repolarization occurred, settling the baseline at 0.88 ± 0.33 mV above the original DC level, while amplitude of reflex responses decreased significantly (Table 2, P < 0.001, paired t-test). Application of the GABA_A_ receptor antagonist, bicuculline (20 µM), depolarized baseline by 2.60 ± 0.75 mV within 4.02 ± 0.49 min (n = 4), which repolarized to baseline (0.16 ± 0.17 mV) after 7.16 ± 1.83 min of continuous application of bicuculline, accompanied by rhythmic bursting (Bracci et al., 1997). Spectral analysis was performed on 5-minute segments of traces recorded in bicuculline at steady-state. Resulting mean RMS value was 8.68 times greater than the RMS calculated from baseline spontaneous activity in control, mirroring the appearance of a rhythmic activity (from 0.10 ± 0.05 in control, to 0.83 ± 0.38 during bicuculline; P < 0.001, paired t-test, n = 11). Also, application of the glycinergic receptor antagonist, strychnine (str, 1 µM), induced bursting activity (Bracci et al., 1996b) with a mean RMS value 2.58 times higher than the one calculated for baseline spontaneous activity in control before str application (from 0.13 ± 0.09 to 0.34 ± 0.2; P = 0.031, paired t-test, n = 5). Finally, when bicuculline (20 µM) and strychnine (1 µM) were co-applied, baseline depolarized on average by 2.97 ± 0.99 mV within 4.11 ± 0.84 minutes from application of drugs and then spontaneously ripolarized to original baseline values within 6.22 ± 0.91 minutes of continuous application of bic + str (n = 12). Appearance of disinhibited bursting (Bracci et al., 1997) corresponds to an 8.46 fold increase in mean RMS compared to pre-drug applications (from 0.12 ± 0.15 mV in control to 1.03 ± 0.27 mV in str + bic; P < 0.001, paired t-test, n = 7). Application of the gap junction blocker, carbenoxolone (CBX, 100 µM), the selective antagonist for adenosine A1 receptors, DPCPX (5 µM), the TRPA1 channel blocker, AP-18 (20, 50, 66 µM), or the TRPV4 channel antagonist, RN1734 (10, 20, 50, 100 µM), did not significantly alter DC levels nor reflex responses. Detailed changes in reflex amplitude, as well as time to peak values and associated statistical analyses, are provided in Table 2.

**Table 2.** Amplitude and time to peak of reflex responses, with statistical comparisons before and after the application of each pharmacological agent.

| Agents | n | Reflex parameter | Before application | After affection of the drug | Test | P value |
| --- | --- | --- | --- | --- | --- | --- |
| CBX | 5 | Amplitude (mv) | 0.33 ± 19 | 0.28 ± 0.16 | paired t-test | 0.180 |
|  | 7 | Time to peak (ms) | 75.82 ± 6.63 | 78.07 ± 4.50 | paired t-test | 0.281 |
| ADO | 6 | Amplitude (mv) | 1.11 ± 0.46 | 0.55 ± 0.26 | paired t-test | 0.003 |
|  | 6 | Time to peak (ms) | 75.67 ± 5.41 | 71.62 ± 3.58 | paired t-test | 0.041 |
| caffeine | 5 | Amplitude (mv) | 0.85 ± 0.47 | 0.80 ± 0.51 | Wilcoxon matched-pairs test | 0.313 |
|  | 5 | Time to peak (ms) | 77.06 ± 5.13 | 80.05 ± 5.10 | paired t-test | 0.039 |
| DPCPX | 5 | Amplitude (mv) | 0.40 ± 0.13 | 0.78 ± 0.68 | Wilcoxon matched-pairs test | 0.813 |
|  | 5 | Time to peak (ms) | 78.36 ± 4.81 | 78.17 ± 5.80 | paired t-test | 0.800 |
| RN1734 | 4 | Amplitude (mv) | 0.42 ± 0.23 | 0.34 ± 0.21 | Wilcoxon matched-pairs test | 0.25 |
|  | 4 | Time to peak (ms) | 79.83 ± 4.05 | 86.68 ± 6.12 | Wilcoxon matched-pairs test | 0.125 |
| Gly | 6 | Amplitude (mv) | 0.62 ± 0.22 | 0.38 ± 0.18 | paired t-test | 0.0003 |
|  | 6 | Time to peak (ms) | 80.93 ± 3.93 | 79.3 ± 2.32 | paired t-test | 0.119 |
| str | 5 | Amplitude (mv) | 0.60 ± 0.17 | 0.90 ± 0.35 | Wilcoxon matched-pairs test | 0.125 |
|  | 5 | Time to peak (ms) | 76.12 ± 6.66 | 82.57 ± 7.78 | paired t-test | 0.101 |

### Pharmacological manipulation of the peak of injury potentials

All pharmachological agents used have been applied in the recording bath in the exact moment when physical injury was produced to the thoracic segment (T10) of the spinal cord. Concentrations, as well as timing of application, were defined as time values before and after the impact (Table 1). DC level of baseline was continuously recorded from VRrL5. The peak of injury-induced depolarization was normalized to that previously elicited by the application of a high potassium medium (10 mM) in the same experiments (Fig. 2A). Among all drugs tested, only RN1734 at concentrations greater than 50 µM significantly reduced, on average, the normalized amplitude of injury potentials by 35.25 % compared to control untreated preparations (P = 0.034, Kruskal-Wallis Test followed by Dunn’s Multiple Comparisons Test; Fig. 2B). However, no significant differences were observed in the onset of injury potentials in response to the application of any of the drugs tested (P = 0.522, Kruskal-Wallis Test followed by Dunn’s Multiple Comparisons Test). In summary, impact-induced depolarization was smaller when RN1734 was perfused to the preparation during impact.

**Figure 2.**
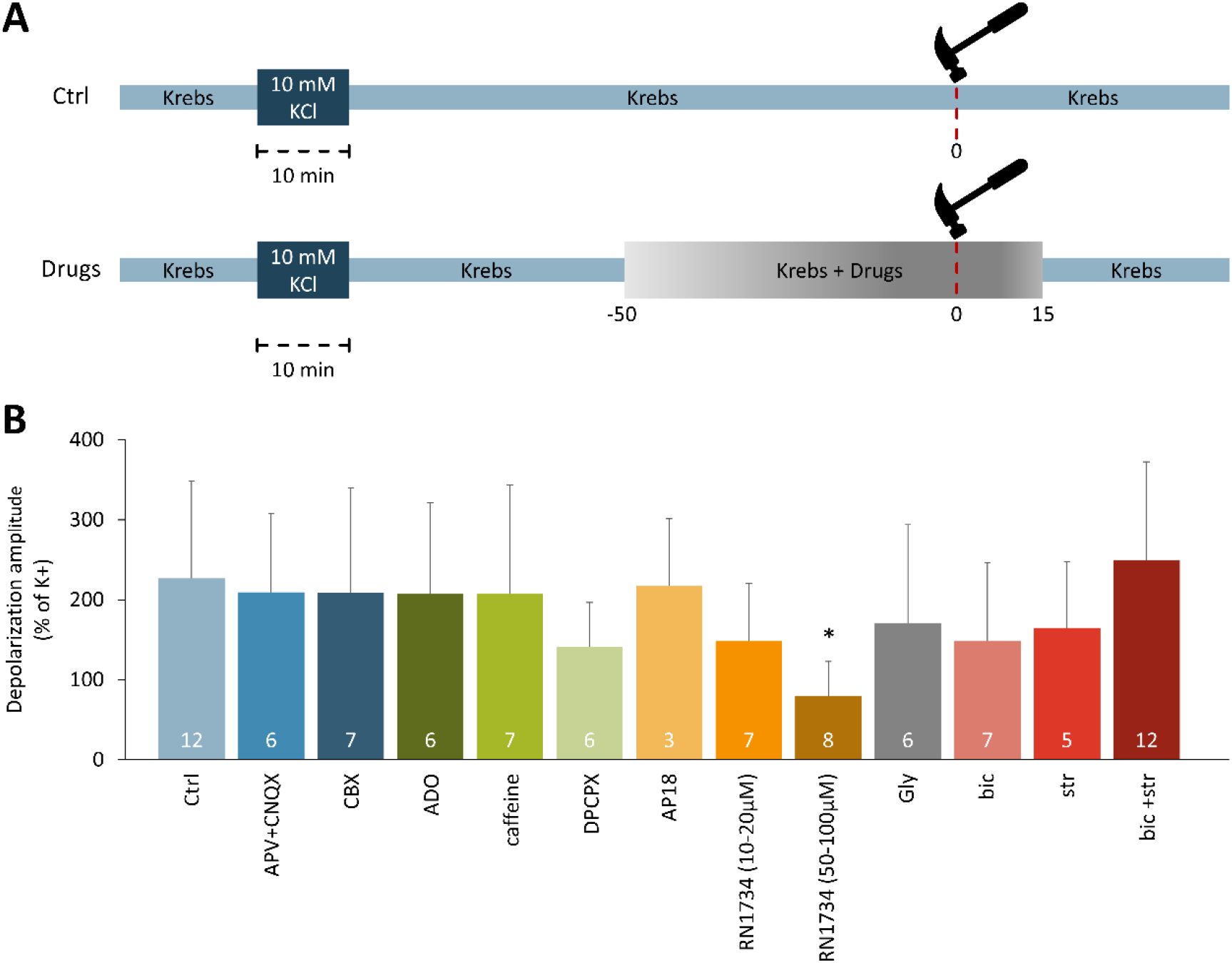
Amplitude of injury-potentials is reduced by TRPV4 antagonism. **A**. Bar chart schematizes the experimental protocols used for impacts on untreated preparations (ctrl, above) and during perfusion with neurochemicals (drugs, below). **B**. Histogram displays the normalized amplitude of injury-induced depolarization recorded from VRrL5 during the impact in the presence of different pharmacological agents. Values are normalized as percentages of potassium-induced depolarizations in the same preparation (* P = 0.034). Total number of experiments in each group is indicated within each bar.

### Pharmacological manipulation of motor reflex responses

Dynamics of the recovery of motor reflex responses following injury varied with each drug tested (Fig. 3A). Amplitude of responses was normalized to pre-impact values for each preparation. Two time-points were considered. In untreated injured preparations, a slow spontaneous recovery in the peak of motor-evoked responses occurred after injury, reaching 28.97 ± 17.17 % of pre-impact after one minute and 45.64 ± 28.05 % after two minutes. Conversely, in preparations treated with CBX, responses partially recovered to 66.39 ± 19.32 % of pre-impact control values immediately after the impact (1 min). At the same time point, RN1734 completely restored pre-impact motor evoked responses (90.70 ± 28.48 %). Both, CBX and RN1734 significantly improved the amplitude of normalized responses one minute after impact, when compared to the peak of normalized motor reflexes in untreated spinal cords (P = 0.004, Kruskal-Wallis Test followed by Dunn’s Multiple Comparisons Test; Fig. 3B). With a longer delay from the trauma (2 min), also bic brought to a full recovery of normalized reflex amplitude values (72.33 ± 16.86 % of pre-impact), (P = 0.040, Kruskal-Wallis Test followed by Dunn’s Multiple Comparisons Test; Fig. 3C). Moreover, no significant difference was observed in the rate of reflex recovery following application of any of the other drugs tested at either time ponts. Collectively, after a physical trauma to the spinal cord, CBX speeded up the early partial recovery of motor evoked responses, while both, RN1734 and bic, stably restored full responses even if with a temporal profile that makes RN1734 more effective than bic.

**Figure 3.**
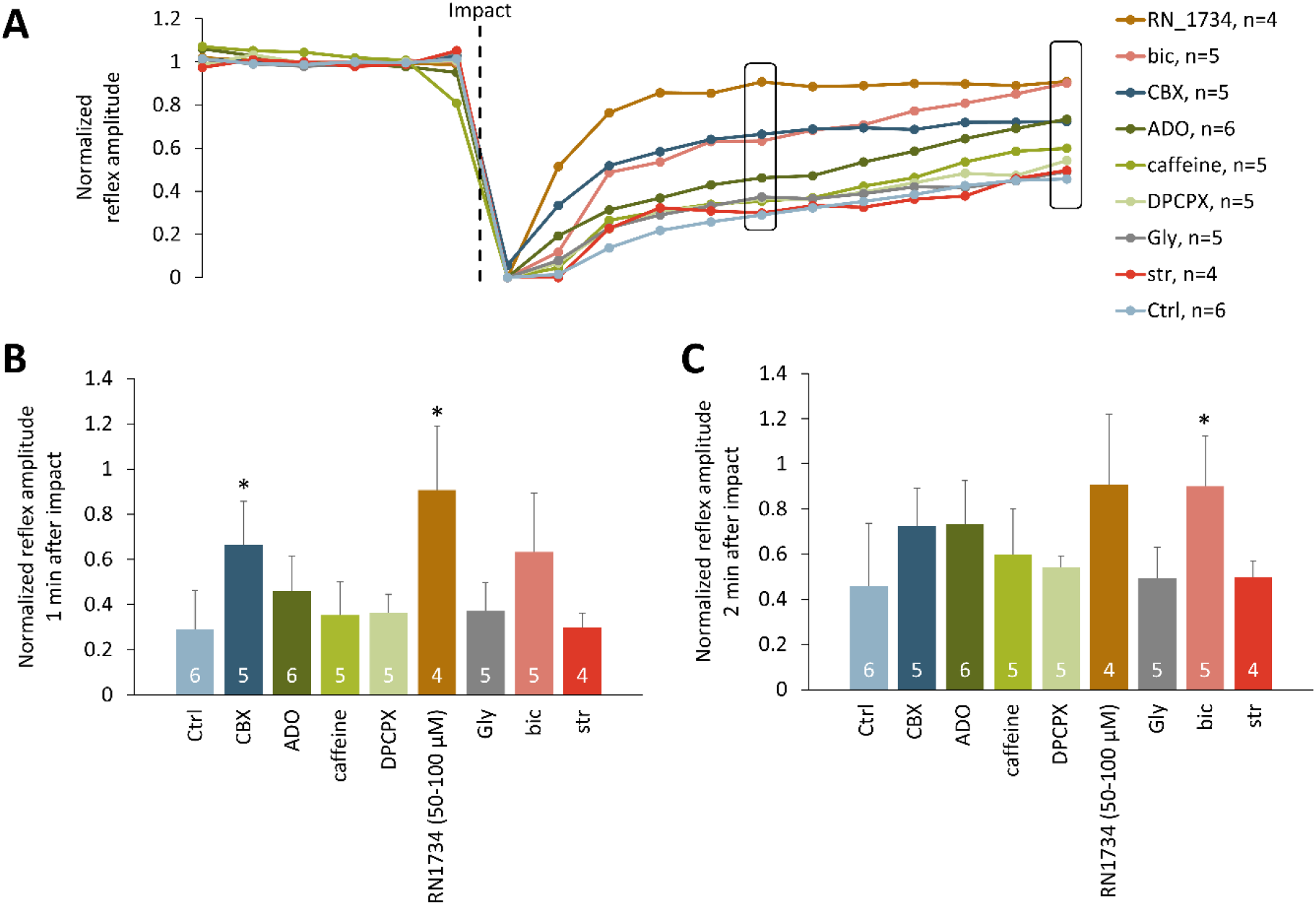
Motor reflex responses after a physical impact to cord recover with different rates depending on drug exposures. **A**. A 3-minute time course representing motor reflex amplitude recorded from VRrL5 after exposure to different drugs. Values are normalized to the average responses before the impact (indicated by the dotted line). **B**. Bar chart illustrates normalized reflex responses one minute after the impact (corresponding to the left square in A, * P = 0.004). **C**. Normalized reflex responses two minutes after the impact (corresponding to the right square in A, * P = 0.040). The total number of experiments in each group is indicated within each bar.

## Acknowledgments

GT is grateful to Mrs. Elisa Ius for her excellent assistance in preparing the manuscript and to John Fischetti for technical support in fabricating the impactor. The study was supported by intramural SISSA grants through the 5xMILLE2020 framework.

## Abbreviations

CNS: central nervous system
Ctrl: control
DAPI, 4’: 6-diamidino-2-phenylindole
DR: dorsal root
DRG: dorsal root ganglia
L: lumbar
P: postnatal
r: right
RMS: root mean square
SCI: spinal cord injury
Th: threshold
T: thoracic
VR: ventral root

## Notes

### Competing Interest Statement

The authors have declared no competing interest.

